# Optimized path planning surpasses human efficiency in cryo-EM imaging

**DOI:** 10.1101/2022.06.17.496614

**Authors:** Yilai Li, Quanfu Fan, Ziping Xu, Emma Rose Lee, John Cohn, Veronique Demers, Ja Young Lee, Lucy Yip, Michael A. Cianfrocco, Seychelle M. Vos

## Abstract

Cryo-electron microscopy (cryo-EM) represents a powerful technology for determining atomic models of biological macromolecules(Kühlbrandt, 2014). Despite this promise, human-guided cryo-EM data collection practices limit the impact of cryo-EM because of a path planning problem: cryo-EM datasets typically represent 2-5% of the total sample area. Here, we address this fundamental problem by formalizing cryo-EM data collection as a path planning optimization from low signal data. Within this framework, we incorporate reinforcement learning (RL) and deep regression to design an algorithm that uses distributed surveying of cryo-EM samples at low magnification to learn optimal cryo-EM data collection policies. Our algorithm - cryoRL - solves the problem of path planning on cryo-EM grids, allowing the algorithm to maximize data quality in a limited time without human intervention. A head-to-head comparison of cryoRL versus human subjects shows that cryoRL performs in the top 10% of test subjects, surpassing the majority of users in collecting high-quality images from the same sample. CryoRL establishes a general framework that will enable human-free cryo-EM data collection to increase the impact of cryo-EM across life sciences research.

## Main

Cryo-EM is one of the fastest-growing areas of structural biology, enabling 3D structure determination of important macromolecular complexes and membrane proteins(Nogales and Scheres, 2015). The impact of cryo-EM ranges from plant biology to human health and disease. Indeed, the widespread growth of cryo-EM has advanced our understanding of pathogens, such as SARS-CoV-2(Rapp et al., 2022), and has had a direct impact on drug development(Wigge et al., 2020).

Cryo-EM is an expensive and highly involved technique. For example, high-end microscopes cost millions of dollars to purchase and require yearly service contracts costing hundreds of thousands of dollars and full-time staff to maintain the instrument. To collect cryo-EM data, users require microscope time for multiple days to collect sufficient high-quality data in addition to waiting weeks or months to access high-end cryo-EM instruments. Thus, increasing the throughput of cryo-EM data collection is critical for advancing biomedical research efforts and reducing the cost of cryo-EM.

Major advances in microscope hardware (e.g., detectors(McMullan et al., 2016) and aberration-free image shift(Cheng et al., 2018; Weis and Hagen, 2020)) have led to rapid data acquisition strategies. Despite collecting hundreds of exposures per hour, much of the collected data is discarded during processing. For example, from 2019 to mid-2021, only 50% of exposures collected in the Cianfrocco laboratory had resolution estimates of < 6 Å. This is because cryo-EM users do not know *a priori* which regions of a cryo-EM grid will produce the highest resolution data. Analysis of micrograph quality across different regions of a grid highlights the complex data landscape users must navigate, where variations of data quality occur locally (within micron-sized areas of squares) and globally (across the entire grid) (**Extended Data Fig. 1**). Cryo-EM samples are challenging samples for data collection, indicating that path planning optimization in addition to rapid data acquisition stands to improve microscope performance and throughput.

Artificial intelligence provides a powerful framework to recognize patterns in complex datasets. Recently, incorporating deep learning models into a reinforcement learning (RL) algorithm enabled RL to surpass professional human performance in challenging games such as AlphaGo(Silver et al., 2016) and Dota 2(OpenAI et al., 2019). These approaches demonstrate superior capabilities in learning strategic moves from simulation data. Given that many problems in the real world involve perception, planning, and decision-making, RL may impact many areas of human life, such as self-driving vehicles(Balaji et al., 2020), robotics(Kalashnikov et al., 2018), healthcare(Yu et al., 2021), and even trading and finance(Hambly et al., 2021). However, the application of RL in life sciences remains underexplored due to the lack of domain knowledge needed to formulate RL into scientific processes.

As a step toward developing fully automated intelligent decision-making for cryo-EM data collection, we present an RL-based framework, cryoRL. The goal of cryoRL is to identify the optimal path across cryo-EM samples to collect as much high-quality data as possible in a given time period (see **Problem Formulation**). Our algorithm combines a low-magnification survey with pretrained classification models that feed into a deep-Q network for learning data collection policies. We benchmarked the performance of cryoRL against human subjects using a cryo-EM data collection simulator, allowing both cryoRL and humans to simulate data collection on the same dataset. Our results show that cryoRL outperforms 9 out of 10 human subjects, suggesting that cryoRL will serve as a framework to automate cryo-EM data collection from all cryo-EM instruments. Our data suggest that optimized path planning with cryoRL will enable an approximately 40% improvement in microscope throughput, thus improving access and throughput on these instruments, thereby reducing the overall cost of cryo-EM.

### Problem formulation

In general, the cryo-EM data collection process can be viewed as a navigation problem in the sample space. The goal is to find a trajectory that maximizes the number of visited holes with high-quality data in a given time. This sequential process involves switching across images at different magnification levels, which can be time-consuming and expensive. We formulate the problem as a Markov decision process (MDP) in the following way: each hole represents a state, and at each state, the agent chooses the next hole to visit from the set of all unvisited holes, which forms the action space. The reward is given by whether the next hole contains good data or not, penalized by the time of moving the microscope stage.

More specifically, our environment is a partially observable MDP because the agent navigates based on the predicted quality scores that are often imperfect. By visiting some holes within a patch (or square), the agent may improve its prediction based on the new information collected locally (*i*.*e*., the true quality scores of the holes visited). This means that the optimal policy depends on the last state and a sequence of states. We, therefore, construct the features for our RL agent with predicted quality scores using observations from multiple previous steps and the summary statistics for the current patches and squares (see **Extended Data Table 1** for details on information used). Note that this also makes it significantly advantageous to apply RL instead of learning-based combinatorial optimization algorithms(Bengio et al., 2021; Mazyavkina et al., 2021), as the latter needs to solve the expensive optimization problem whenever a model is updated.

An effective planning algorithm, thus, should be able to identify promising regions with plentiful high-quality holes and program the microscope to focus on those regions to avoid frequent image switching. To achieve this, cryoRL aims to optimize an objective function as follows:

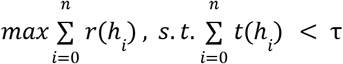

where *τ* is a prespecified time duration, and *r*(*h*_*i*_) is a reward for visiting a hole *h*_*i*_ based on the quality of *h*_*i*_ as well as the time cost *t*(*h*_*i*_) of the microscope movement (see section *Time cost and RL Rewards*). In reality, the hole quality predictions can be noisy, posing additional challenges.

#### Justification & formulation for reinforcement learning (RL) in cryo-EM data collection

We propose to solve such an optimization problem by reinforcement learning (RL), which is well suited for sequential decision-making tasks. In the literature, the decision-making problem is formulated either as a multi-armed bandit (MAB) or RL. MAB makes the decision solely based on the quality of the arm, which does not provide effective path planning for our problem. As described above, our problem can be formulated as an MDP, so RL is an appropriate technique for our problem. Thus, we argue that RL is more suitable for our problem than other approaches, such as learning-based combinatorial optimization(Bengio et al., 2021; Mazyavkina et al., 2021).

For cryo-EM data collection, we need to use predicted quality during path planning. Combinatorial optimization techniques such as genetic algorithms and simulated annealing perform path planning based on prediction and do not consider observations. Thus, they usually do not perform well when the prediction is poor. Compared to combinatorial optimization, RL can incorporate observations into path planning for better decision-making. For instance, when there are too many low-quality holes observed in the current square, a good policy learned in cryoRL will choose to switch to another patch or grid for further exploration, even if the prediction indicates that the square has ‘good’ holes.

To further justify the use of RL, we argue that: 1) our RL features build on both the predicted score of a single hole with relatively low quality and global features reflecting the quality of patches and squares. Thus, it has the potential to do better than combinatorial optimization, which is based solely on local information. 2) During the test phase, new labels will be collected for the visited holes, from which the policy can improve itself by updating the global features. A combinatorial optimization algorithm may not enjoy such benefits.

RL is generally less heuristic than other optimization solvers such as Genetic Algorithm(Weise, 2009) and Simulated Annealing(Kirkpatrick, 1984). In related work, we compared the outcome of RL with the outcomes of GA and SA and found the performance of RL to be significantly better(Fan et al., 2022). Furthermore, although RL is computationally expensive in the training phase, it is quick when we deploy the trained policy within the system. Thus, it can make real-time decisions.

#### Time cost and RL Rewards

In real cryo-EM data collection, the time needed to collect a micrograph varies depending on how far the stage needs to be moved from its previous position. In this study, we defined the time cost *t*(*h*_*i*_) as in **Extended Data Table 2**. We combine the time cost *t*(*h*_*i*_) with the score of micrographs, where ‘good’ micrographs have high-resolution information present as measured by the maximum CTF-estimated resolution (“CTFMaxRes”). While CTFMaxRes is not a direct measure of particle quality in the micrographs, it is an approximate assessment of sample density and ice thickness. We use CTFMaxRes in this work, given its widespread use as a data curation metric, but, importantly, our framework can generalize to any metric for optimizing RL rewards.

When the CTFMaxRes of the micrograph collected from *h*_*i*_ is lower than a given threshold *T*, the reward for each hole visit *r*(*h*_*i*_) is defined as:

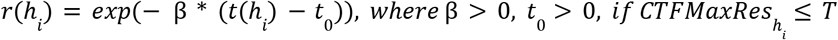

In this study, we assigned *β* = 0. 1, *t*_0_ = 1. 0, and *T* = 6 Å (unless specified otherwise). When the CTFMaxRes of the micrograph collected from *h*_*i*_ is larger than the given threshold *T, r*(*h*_*i*_) is defined as:

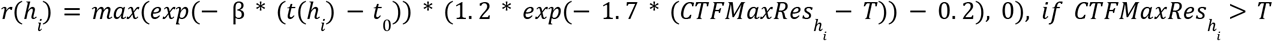

Thus, a partial reward can be gained if the obtained CTFMaxRes is only slightly higher than the threshold *T*. In this study, we presented data with *T* = 6 Å.

#### Snowball baseline policy

We designed a snowball baseline to compare with the results of cryoRL. This baseline estimates the quality of all the candidate holes in the dataset and then will start from a random hole target and aims to maximize the total number of micrographs in a given time duration regardless of the quality. To train this policy, we used *T* = 30 Å in the reward function so that the reward only depends on the time cost.

#### “Predict-Rank” policy

We designed another policy, “Predict-Rank,” as a heuristic baseline to compare with the results of cryoRL. This “Predict-Rank’’ policy used the predicted quality scores and ranked the grid-level images based on their overall predicted quality. For simplicity, we labeled holes with predicted CTFMaxRes below 6 Å as “good.” This is followed by a secondary ranking on the square images and is then followed by a third ranking on the patch-level images. This policy started from a random position and then moved to the “best patch” in the “best square” in the “best grid area,” collected all the predicted good holes, and then moved to the “second best patch” image, etc. This “Predict-Rank” policy serves as a very strong baseline when the prediction of the regressor is accurate.

### Optimization of path planning on cryo-EM samples with cryoRL

Given an unexplored cryo-EM grid, the goal of a data collection session is to obtain the maximal number of micrographs with high-resolution information in a limited time. Importantly, the data collection path needs to be optimized by reducing the time cost associated with stage movement and Z-height adjustment of the microscope. Therefore, two critical problems in cryo-EM data acquisition are 1) quality prediction at the low or medium magnification and 2) trajectory planning during the data collection session.

In traditional cryo-EM data collection, users utilize a sequential approach to interrogate, assess, and steer data collection (**Fig. 1A**). First, a user will generate an atlas of low magnification “grid-level” images to obtain an overview of the sample (**Fig. 1A**). The user will examine the atlas and choose to collect micrographs from the holes in one patch, from a single square. Specifically, for each square selected, the user will capture a “square-level” image and several “patch-level” images at higher magnifications to visualize the overall shapes and relative intensities of each hole (**Fig. 1A**). Depending on the quality of the collected micrographs, the user decides to collect in the same patch, a different patch, or a different square. This iterative process will continue until the user finds a pattern that produces the highest quality data.

**Figure 1.**
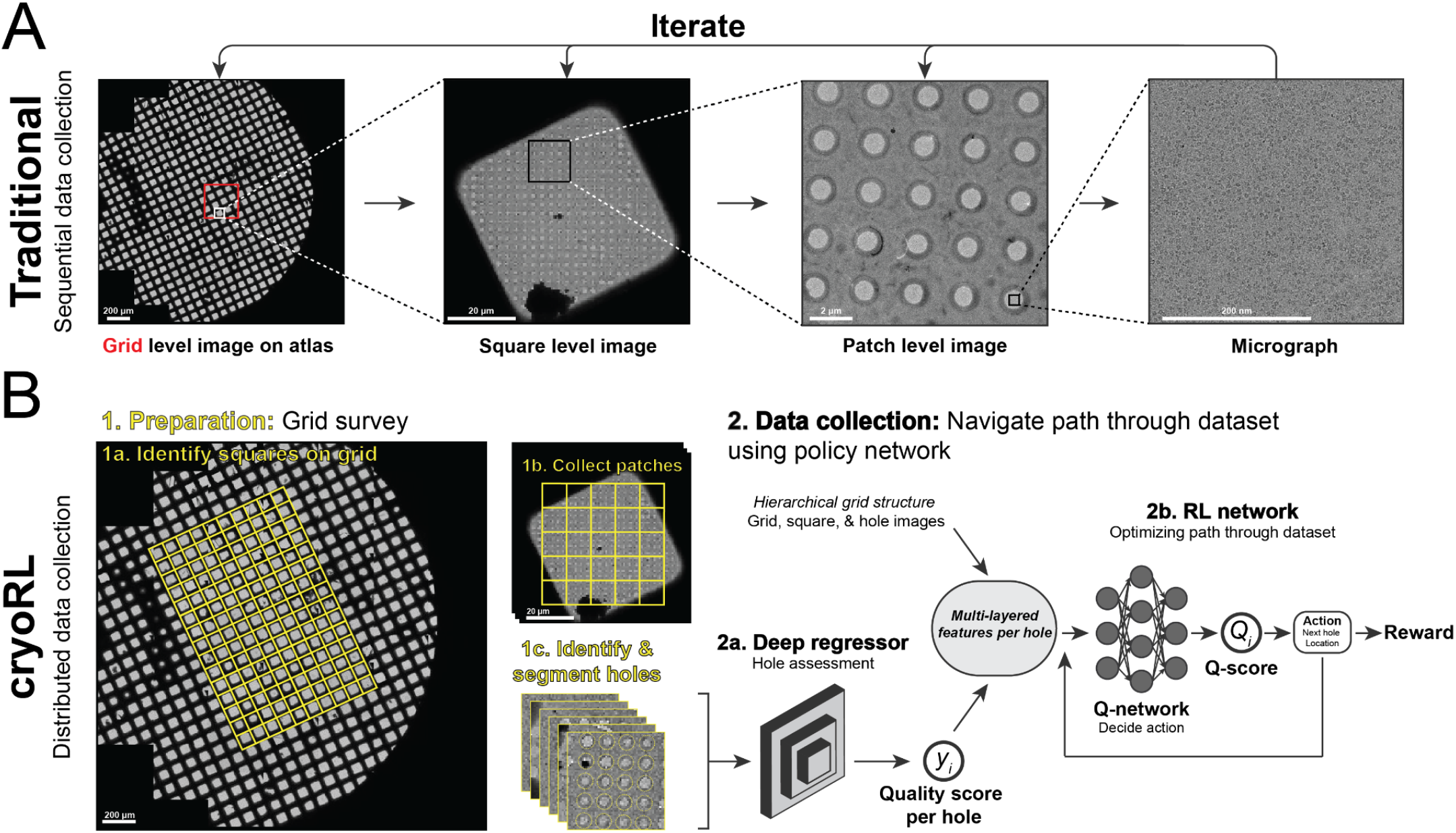
Overview of cryoRL approach. (A) Traditional data collection involves a sequential investigation of a cryo-EM sample atlas, squares, and holes found within patches to obtain high-magnification micrographs. (B) Overview of cryoRL approach. (1) Preparation: Low magnification survey of square- and patch-level images are collected on an area of interest. (2) Data collection: A deep Q-network is trained using hole assessment (“Deep Regressor”) and grid image hierarchies are inputs. The network learns to output an action that is based on the current state on the grid.

To automate cryo-EM data collection, we designed an RL-based data collection algorithm: ‘cryoRL’ (see **Problem Formulation**). CryoRL aims to optimize cryo-EM data collection by combining supervised regression and reinforcement learning (**Fig. 1B**). First, a deep regressor will predict a quality score for each hole image at medium magnification. Subsequently, an RL model will use these quality scores and the hierarchical information on the grid to output a trajectory of actions, i.e., to decide on the fly which positions should be used for collecting high-resolution micrographs.

In this work, we define micrograph quality based on the amount of signal in Fourier space for a given micrograph. Specifically, we calculate the maximum resolution at which the contrast transfer function (CTF) can be detected. This value - “CTFMaxRes” - is a routine value that is a standard part of cryo-EM processing(Cianfrocco and Kellogg, 2020; Rohou and Grigorieff, 2015). CTFMaxRes is a valid metric considering that it correlates with particle number and the signal-to-noise ratio of the images. Importantly, CTFMaxRes is a widely used parameter to filter out “bad” micrographs during single particle processing, such as removing micrographs with a CTFMaxRes of > 6 Å. In cryoRL, the CTFMaxRes value of a micrograph defines the quality score of the corresponding hole-level image.

CryoRL relies on low-magnification grid mapping to facilitate path planning across the grid. We call this a “distributed” approach for data collection, which is contrary to the traditional “sequential” data collection strategy (**Fig. 1**). A distributed view of a grid provides a landscape of the grid, which is essential for trajectory planning. After collecting patch-level images across a square, which can take 1-2 minutes per square, all holes will be automatically identified and segmented. These hole images will constitute a candidate pool in cryoRL. Since these images are low magnification, this is automated and does not constitute a time-consuming step.

The reward function is a critical aspect of RL implementations as these rewards will guide RL behavior(Li, 2017). In cryoRL, we developed a reward scheme that values finding good micrographs from neighboring holes, where finding a good micrograph within the same patch has the highest reward (see **Problem Formulation** & **Methods**). The reward function also reflects realistic considerations when operating the instrument: moving to neighboring holes is faster than switching areas of a square or region of a grid. cryoRL will reward the action of taking a “good” micrograph, and the reward amount is negatively correlated to the time it takes to perform the action (**Extended Data Table 1**, see **Methods**). Therefore, cryoRL takes account of both quality prediction and trajectory planning in a real-world setting with a practical consideration of the time cost. For a baseline comparison, we designed a “snowball baseline,” which aimed to maximize the total number of micrographs collected regardless of the quality, to compare the results of cryoRL with the outcomes of a semi-random data collection policy (see **Problem Formulation** & **Methods**).

To test and implement cryoRL, we performed all path planning offline by running cryoRL on systematically collected datasets (see **Methods**). For these datasets, we imaged every hole across many squares (> 10) to provide cryoRL with a landscape of holes from which to choose.

### CryoRL effectively navigates cryo-EM samples to identify high-quality data

We tested the performance of cryoRL offline on a cryo-EM sample of rabbit muscle Aldolase on a Quantifoil 1.2/1.3 grid from which we collected a systematic dataset (**Fig. 2A**, see **Methods**). For this sample, we trained the deep regressor and deep Q-network on a subset of the cryo-EM grid. The deep regressor successfully captured the correlation between hole images and CTFMaxRes values (**Extended Data Fig. 2A & 3A**). Using these trained models, cryoRL collected micrographs from “good” holes on the grid (**Fig. 2B & 2C**). Given that the overall candidate holes in this dataset are low quality (less than 40% of candidates had a CTFMaxRes < 6 Å (**Fig. 2B**) - cryoRL collected ∼80% of images with a CTFMaxRes < 6 Å (**Fig. 2C**). CryoRL also outperformed the Predict-Rank policy, which uses the information of the quality prediction but not reinforcement learning-based path planning (**Fig. 2C**).

**Figure 2.**
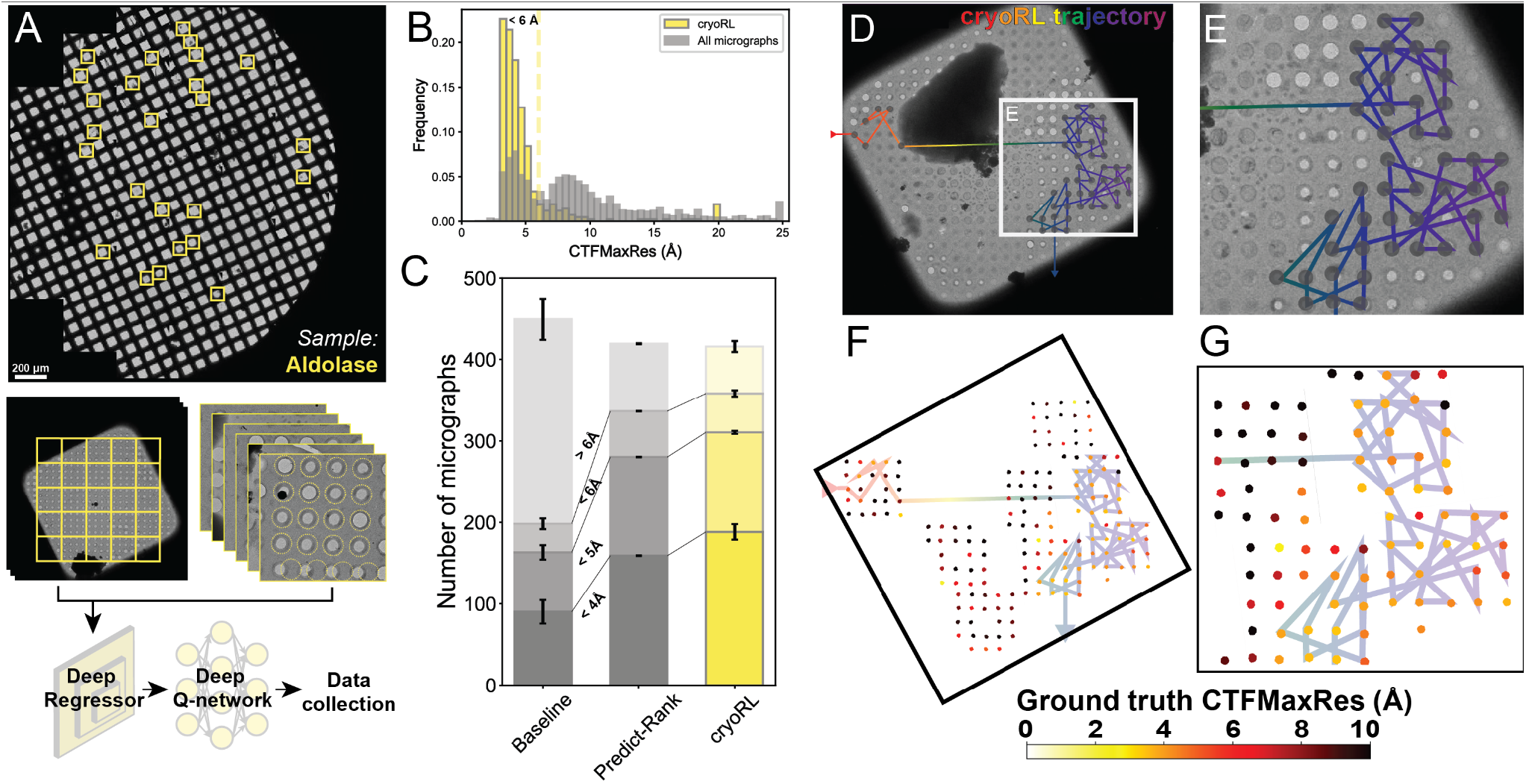
cryoRL collects high quality cryo-EM data. (A) Overview of cryo-EM grid used for systematic data collect. Yellow squares indicate regions where almost every hole was collected. (B) Comparison of CTFMaxRes value for all micrographs (gray) vs. cryoRL (yellow). (C) Comparison of cryoRL, Predict-Rank and snowball baseline. (D) Example square showing path of cryoRL. Holes were shaded if they were in the trajectory of cryoRL. (E) Zoom-in on panel in (D). (F) & (G) cryoRL trajectory overlaid on ground-truth CTF values per hole.

To visualize the behavior of cryoRL, we overlaid a cryoRL trajectory on the cryo-EM sample (**Fig. 2D-2G**). cryoRL visited six squares during data collection. Zooming in on a selected square, we see that this specific square contained holes with different ice thicknesses and quality (**Fig. 2D**). CryoRL successfully avoided empty holes or holes with very thick ice (**Fig. 2E**). Compared to the ground truth CTFMaxRes of all the candidate holes, we found that cryoRL collected micrographs with low CTFMaxRes only in the patches with many high-quality holes (**Fig. 2F & 2G**). Therefore, cryoRL learned an effective policy that minimizes the time cost increase from moving to a different patch. We conclude from these results that cryoRL successfully collected high-quality micrographs when trained on a small subset of the dataset.

### Transferred models enable effective path planning on different cryo-EM samples and grid types

Our result on Aldolase showed that cryoRL learns a policy to collect high-quality micrographs when both the regressor and RL network are trained from a subset of the same grid. However, RL models need to be trained on a subset of the dataset where all the possible holes are collected with a ground truth CTFMaxRes. A dataset like this with a sufficient sample size for training is usually unavailable and not practical to obtain. Therefore, RL models must have good transferability so they do not need to be trained on the same dataset.

To test the transferability of RL models, we collected a dataset of a different cryo-EM sample, Apoferritin (for transferability comparisons, see **Extended Data Table 3**). We utilized a general deep regressor that was trained on 100,578 hole images from the Cianfrocco laboratory from a variety of sample types with associated CTFMaxRes values (see **Methods, Extended Data Fig. 2B & 3B**) in combination with the deep-Q network from Aldolase (**Fig. 3A**). The resulting images collected by cryoRL showed a strong enrichment for images < 6 Å (**Fig. 3B**), indicating that the transferred models’ allowed high-quality data collection. The overall performance of cryoRL is 1.5 times above the baseline with ∼80% of collected images < 6 Å (**Fig. 3C**). Surprisingly, even though the performance of the regressor was not perfect (**Extended Data Fig. 2B & 3B**), the Predict-Rank policy performed similarly to cryoRL (**Fig. 3C**).

**Figure 3.**
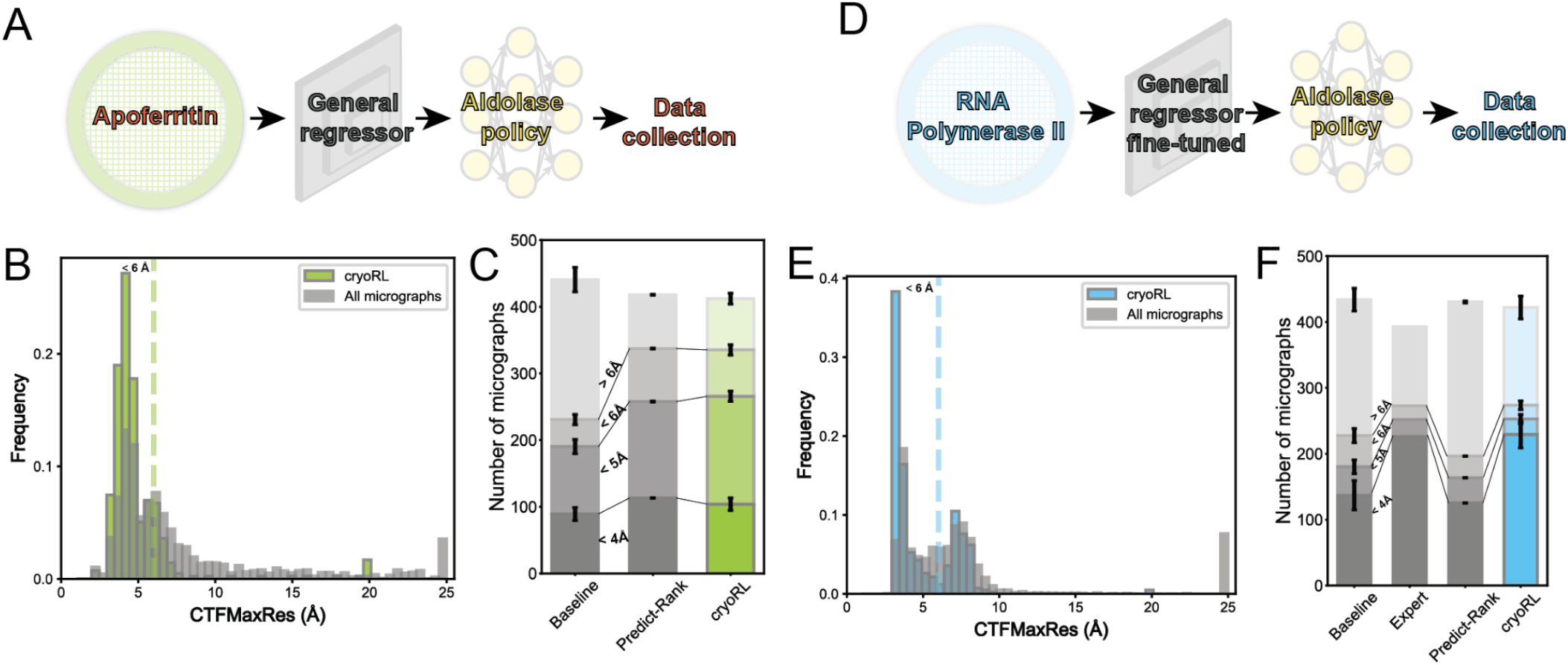
Transferred models enable effective data collection by cryoRL. (A) Schematic for apoferritin dataset. (B) CTFMaxRes histogram for cryoRL vs. all images. (C) Overall performance of cryoRL, Predict-Rank, and snowball baseline. (D) Schematic for RNA polymerase II (RNAPII) dataset. (E) CTFMaxRes histogram for cryoRL vs. all images. (F) Overall performance of cryoRL, Predict-Rank, and snowball baseline.

Given the performance of cryoRL on Apoferritin, we switched to a different sample on a different grid: RNA Polymerase II (RNAPII) on Au-Flat 1.2/1.3. First, we carefully divided the atlas into halves. An expert (Michael Cianfrocco) freely collected on one half of the grid, while the other half was used for the systematic data collection that served as the ground truth for cryoRL. CryoRL utilized a deep regressor fine-tuned from the pretrained general regressor with a subset of holes from the RNAPII grid (see **Methods, Extended Data Fig. 2C & 3C**). For the deep-Q network, instead of training on the same grid, we used the deep-Q model from the Aldolase dataset. Using the fine-tuned deep regressor and transferred deep-Q network (**Fig. 3D**), cryoRL collected high-quality images from the RNAPII dataset (**Fig. 3E**). A comparison of cryoRL versus the baseline shows two times more images below 4 Å (**Fig. 3F**). On the other hand, the Predict-Rank policy performed similar to the baseline (**Fig. 3F**), showing the inconsistency when the prediction result is noisy. For the data collected by the expert, we translated the trajectory with the time costs used in this study (**Extended Data Table 1**) and truncated the trajectories at 480 min. The performance of the expert was similar to cryoRL (**Fig. 3F**).

### CryoRL outperforms humans

To test human performance against cryoRL on the same dataset, we designed and implemented a cryo-EM simulator (see **Methods, Supplemental Movie 1**). We used a cryo-EM simulator to ensure that both human users and cryoRL have access to the exact same grid, square, hole, and exposure data to ensure an accurate comparison. We compared the performance of cryoRL versus human subjects on a new systematic dataset of rabbit muscle Aldolase on a different grid type: UltrAufoil 1.2/1.3 (“Aldolase^Au^”, **Fig. 4A**). This dataset consisted of 5822 hole images and their ground truth CTFMaxRes from 30 squares. We asked ten users ranging from 3 months to 10 years of cryo-EM experience (see **Methods**) to collect data with the simulator as if they were in a real data collection session. Furthermore, similar to the RNAPII dataset, we also divided the atlas into halves and assigned one half for expert data collection.

**Figure 4.**
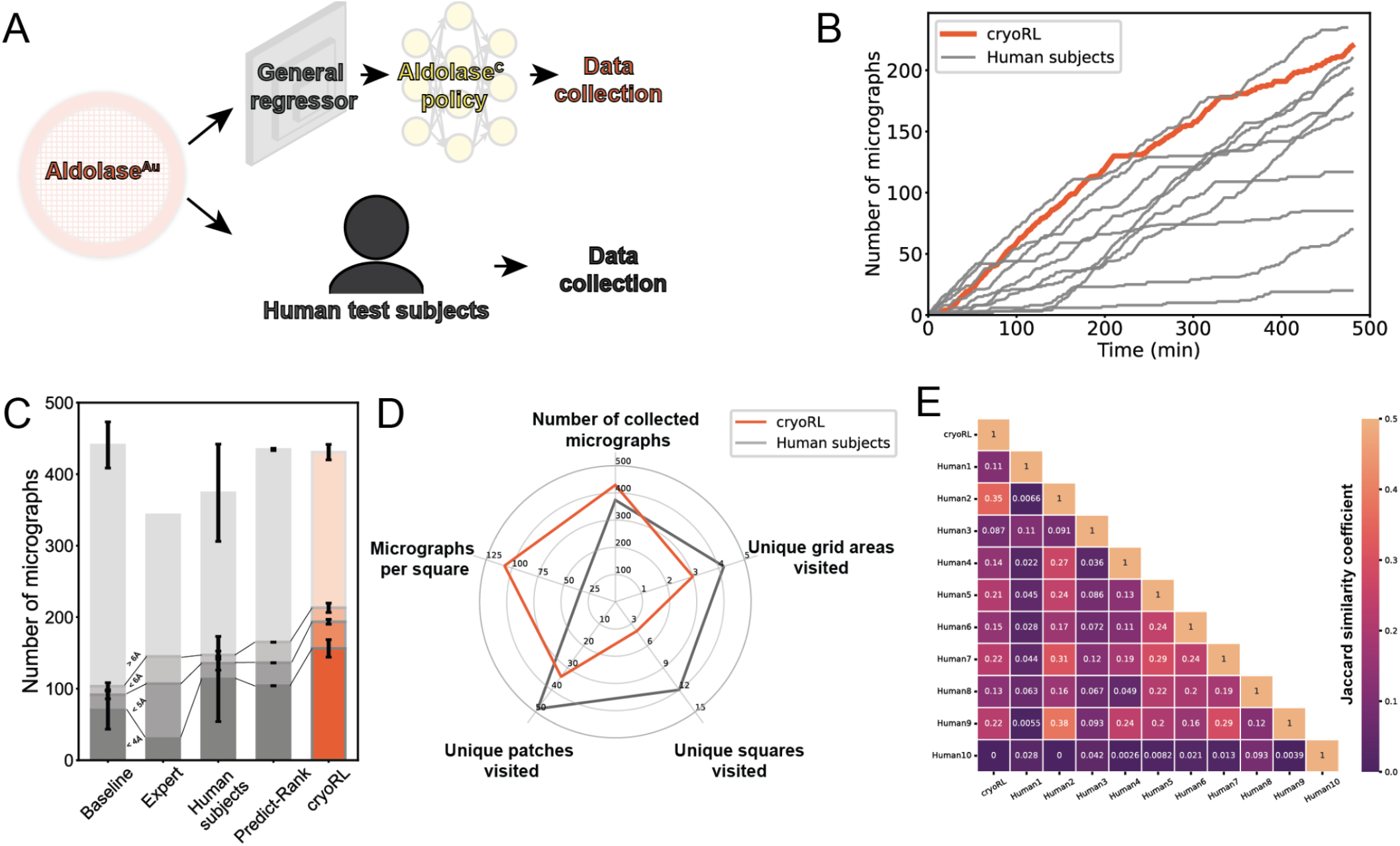
CryoRL outperforms humans. (A) Schematic for Aldolase^Au^ dataset. (B) Number of micrographs with CTFMaxRes < 6 Å over the data collection time for cryoRL and human test subjects. (C) Overall performance of cryoRL, Predict-Rank, snowball baseline and human test subjects. (D) Statistics of data collection behaviors of cryoRL and human test subjects. (E) Pairwise Jaccard similarity coefficients between the micrographs collected by cryoRL and human test subjects.

We compared the performance of human subjects with cryoRL that used the pretrained general deep regressor (**Extended Data Fig. 2D & 3D**) and the DQN trained with Aldolase^Au^ (**Fig. 4A**). From the trajectories of the number of micrographs with CTFMaxRes < 6 Å over time, we found that cryoRL outperformed 9 out of 10 human test subjects (**Fig. 4B**). Interestingly, human subjects showed a large variance in performance. The slopes of the lines indicated the “good data rate.” CryoRL showed a consistent “good data rate,” whereas the rate of most test subjects varied over time (**Fig. 4B**).

We quantified the average performance of human test subjects and the expert and compared them with that of the baseline, the Predict-Rank policy, and cryoRL (**Fig. 4C**). The baseline showed that this dataset contained only 12% of targets with a CTFMaxRes < 6 Å (**Fig. 4C**). Surprisingly, the performance of the expert is even worse than the baseline (**Fig. 4C**). The average performance of humans exceeded the baseline, but the variance was significant (**Fig. 4C, Extended Data Fig. 4**). Predict-Rank policy performed similarly to the average human test subjects (**Fig. 4C**). CryoRL outperformed humans by about 43% on average, with a much lower variance, suggesting higher consistency (**Fig. 4C**).

Next, we wanted to determine the behaviors of cryoRL vs. human subjects. We found that cryoRL only visited four unique squares. In contrast, human users, on average, visited 12 squares (**Fig. 4D**). Consequently, cryoRL collected over 100 micrographs per square, three times more than the human test subjects (**Fig. 4D**). Given this behavior, we conclude that in minimizing stage movements, cryoRL collected more micrographs in total and more high-quality micrographs as a result.

We wanted to compare if the good micrographs collected by cryoRL and human subjects overlap. To do this, we quantified the pairwise similarities between the micrographs in each trajectory with the Jaccard similarity coefficient (**Fig. 4E**). Surprisingly, we found that human subjects had an average of 0.13 similarity coefficient (0 is uncorrelated, 1 perfectly correlated), further underscoring the variability of data collection (**Fig. 4E**). However, the three users who were able to collect over 200 micrographs < 6 Å (Human-2, -7, and -9) had the highest similarity to cryoRL (**Extended Data Fig. 4, Fig. 4E**). Not surprisingly, the similarity among the best-performing users was relatively high (**Fig. 4E**). On the other hand, the three users who collected less than 100 good micrographs (Human-1, -3, and -10) had weak concordance with cryoRL and among themselves (**Extended Data Fig. 4, Fig. 4E**). This comparative analysis showed that good micrograph selections were alike within this dataset, but every bad micrograph selection was bad in its own way.

Finally, to show that the data collected by cryoRL on this dataset was more consistent than human test subjects, we processed the micrographs collected by cryoRL and human test subjects in the same way (**Fig. 5A**). Since cryoRL started from a random position we sampled ten trajectories out of the 50 trajectory runs produced by cryoRL. The final structure from the datasets cryoRL collected had resolutions ranging from 3.8 Å to 4.3 Å (**Fig. 5B & 5C, Extended Data Fig 5**). In contrast, the resolutions of the structures from the datasets human test subjects collected ranged from 3.9 Å to 9.0 Å ((**Fig. 5B & 5C, Extended Data Fig 5**). Comparing the results from these human test subjects versus cryoRL shows that cryoRL archives consistency that human subjects could not achieve.

**Figure 5.**
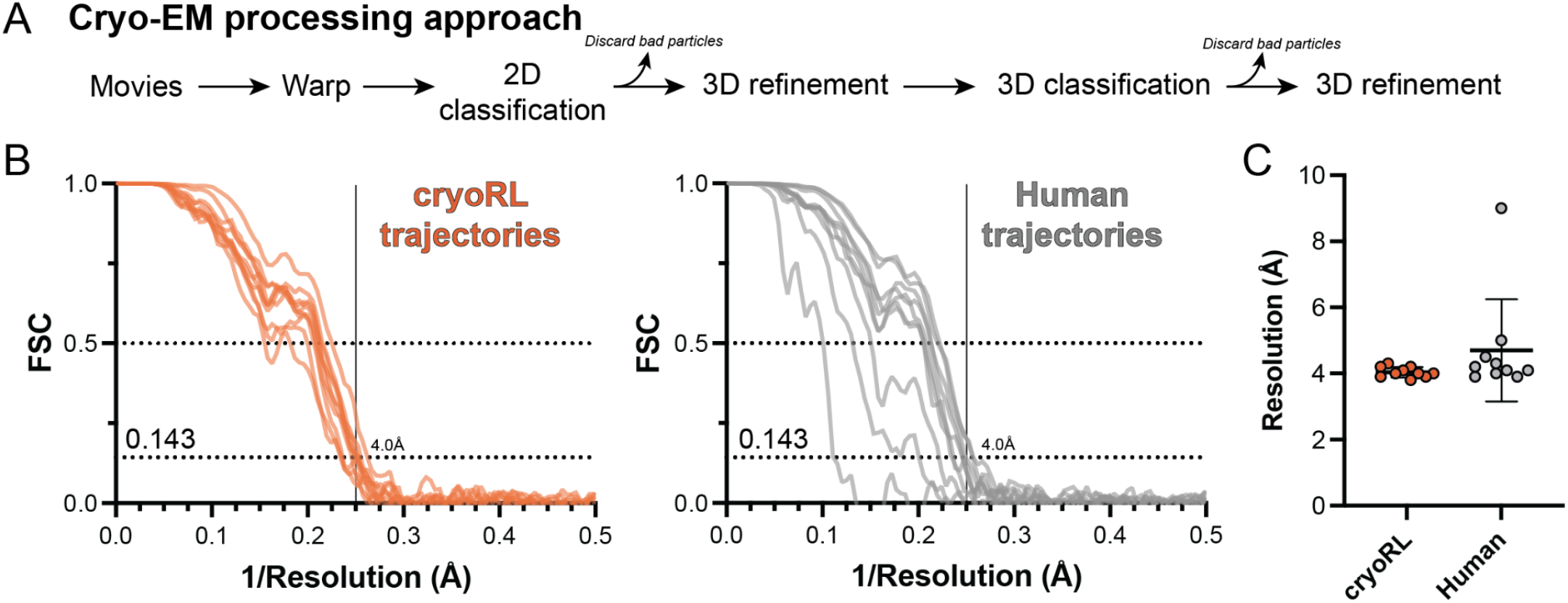
CryoRL data collection trajectories produce consistent reconstruction output. (A) Standard workflow for analysis of all cryoRL and human datasets from Aldolase^Au^ simulator data. Fourier shell correlation curves (FSC) for ten cryoRL (left) or human (right) trajectories after analysis using workflow in (A). (C) Comparison of FSC@0.143 resolution between cryoRL and human final reconstructions from trajectories. Error bars show S.D.

## Discussion

In this work, we show that cryoRL optimizes the path of cryo-EM image collection to maximize high-quality images in a limited time across four independent datasets. Among our four experiments, three were tested with transferred models, showing the generalizability of cryoRL. Our results showed that the RL networks can provide robustness and place cryoRL in the top 10% of human study subjects. Importantly, the deep-Q network is highly transferable as all four experiments shared the same deep-Q model (**Extended Data Table 3**).

Our design of low-magnification prediction with RL policy training enables cryoRL to identify optimal paths across cryo-EM samples. The deep regressor provides a consistent prediction of the data quality at the hole level, allowing cryoRL to map thousands of holes, a task that is nearly impossible for human users. We believe the challenge of mapping paths is highlighted in the cryo-EM simulator: many human subjects cannot keep a consistent rate of data quality collection (**Fig. 4B**).

CryoRL outperforms average cryo-EM users due to consistent data collection rates on local regions of high-quality images. Analysis of cryoRL behavior versus human subjects showed that cryoRL visited four squares, whereas human subjects visited an average of 12 squares (**Fig. 4D**). Moreover, from these four squares, cryoRL obtained a near-constant “good data collection rate” (**Fig. 4B**), whereas human subjects showed large variability in the rate of collection (**Fig. 4B**). From these data, we conclude that cryoRL can consistently collect high-quality data.

Data quality is difficult to infer from hole images since their magnification is usually not high enough. The Predict-Rank policy is a heuristic baseline that only depends on the result from the predicted CTFMaxRes. We found that the performance of Predict-Rank is very unstable throughout our four experiments. However, reinforcement learning was able to provide robustness when the regressor result was noisy.

CTFMaxRes is a metric that has been used for years in the field as a data curation approach. However, we acknowledge that, in many cases, it does not reflect the true quality of a micrograph. The generality of the cryoRL framework allows for any micrograph metric to serve as a target for reward. The deep regressor can also be trained to predict other metrics, like data quality measurement output by MicAssess(Li et al., 2020) or picked particle numbers, as long as the training can converge.

The reward table within cryoRL will enable tuning cryoRL for different data collection behaviors. For example, by increasing the reward for images from different squares, we can encourage cryoRL to explore larger regions of a sample, as in data screening. The time costs and the rewards associated with them can also be tuned to reflect the specifications of different instruments. We believe that cryoRL has great flexibility and potential to automate both data collection and data screening in various scenarios.

### Limitations

Although we showed that cryoRL collected data better than most of the human test subjects, there are some limitations of our framework. First, the performance of cryoRL is directly related to the accuracy of the deep regressor. Although the RL networks provide some robustness to imperfect quality prediction results, if the grid type is not present in the training data, the performance of a general model will not be very accurate. For example, in the RNAPII dataset, the grid type was not present in the training data of our general model. We thus had to fine-tune the general regressor with a subset of the RNAPII data. As transferability is one of the biggest hurdles in the machine learning field, training a general model that will always be accurate on most datasets will be challenging. On the other hand, given that some samples may have specific hole types or may be visible at hole magnification (e.g., viruses, microtubules), a sample-specific deep regressor could be incorporated into cryoRL for increased accuracy of prediction. In the future, we anticipate that the regressor may also learn from the data just collected so that the decision-making can be improved over time. Semi-supervised learning(Ouali et al., 2020) or active learning(Ren et al., 2021) may be useful techniques that can be applied to cryoRL in the future. An online model will be ideal.

The “distributed” data collection scheme adds an up-front time cost to cryo-EM imaging sessions. In the current framework, the systematic survey may collect images that are clearly empty or too thick. We believe that the survey can be more efficient using human or AI-assisted square selection to identify the most promising squares to start(Kim et al., 2021). Also, we believe that z-height adjustment can be skipped when collecting patch images since they are merely used to provide the candidate hole images, so taking these patch images only takes 1-2 min per square.

CryoRL is proposed as an algorithm for optimized path planning on the cryo-EM grid, and it is not available in the current data collection software yet. Also, cryoRL does not include any square or hole finders; therefore, other algorithms (Bouvette et al., 2021; Cheng et al., 2023; Kim et al., 2021) are needed to be used in combination with cryoRL. Moreover, the square and hole detectors and square classifiers provide an automatic solution for collecting the prerequisite information for path planning in cryoRL.

Overall, we believe that optimized path planning using approaches like cryoRL will allow efficient microscope and expert-level performance without the need for human intervention. From our results, we estimate that microscopes will be 140% more productive than current usage, thus expanding the pool of available microscope time for users worldwide on high-end cryo-EM instruments. This will greatly increase the throughput of cryo-EM projects and help to reduce labor costs, and increase access to microscopes.

## Acknowledgments

We thank members of the Vos and Cianfrocco laboratories, especially Hye Jee Hahn and Nicholas Vangos for preparing cryo-EM samples of aldolase and apoferritin; Christopher Lilienthal for help with Leginon/Appion database interrogation. We thank the Protex facility at the University of Leicester, Louise Fairall, and Christos Savva for mouse apoferritin plasmid and protocol for purification. We thank Lucas Farnung for preparing the RNAPII samples. We thank Liz Kellogg, Joey Davis, and Bolei Zhou for giving us comments on this manuscript. The research reported in this publication was supported by the University of Michigan Cryo-EM Facility (U-M Cryo-EM). U-M Cryo-EM is grateful for support from the U-M Life Sciences Institute and the U-M Biosciences Initiative. This work was supported by R01GM143805 and S10OD020011 (M.A.C. & Y.L.) in addition to DP2-GM146254 (S.M.V).

## Author Contributions

Y.L., Q.F., J.C., M.A.C., and S.M.V conceived the project. M.A.C. and S.M.V. co-led the project. Y.L. and M.A.C. collected systematic datasets. Q.F. developed the reinforcement learning framework. Y.L. and Q.F. incorporated reinforcement learning into the cryo-EM data collection scheme and performed learning experiments. Y.L. developed and ran data analysis. V.D., J.Y.L, and L.Y. developed the cryo-EM data simulator. E.R.L. processed the apoferritin datasets. Y.L., Q.F., Z.X., M.A.C., and S.M.V. wrote the manuscript.

## Data availability statement

The source code of cryoRL is can be found on GitHub: https://github.com/IBM/CryoRL-pytorch. The cropped hole images for “Aldolase” can be downloaded from https://github.com/yilaili/cryoRL-pytorch-data. The full database files and images are hosted on a Globus endpoint in the Cianfrocco laboratory.

## Methods

### Deep regressors

We applied deep regression to estimate the CTFMaxRes of each hole, which has a maximum value of 20 Å. The regressor, backboned by ResNet-50, takes as the input the holes cropped out from hole-level images based on the location information provided in the metadata. It is trained for 50 epochs with the Adam optimizer(Kingma and Ba, 2014) using *l*_2_ loss unless specified otherwise. A cosine scheduler(Loshchilov and Hutter, 2016) is adopted with an initial learning rate. In addition to the implementation in this study, the quality of a hole can also be categorized by a classifier, as done in(Fan et al., 2022). However, the classifier has to be re-trained when the categorization criteria change.

The same model architecture was used for the training of the regressor for Aldolase, the general regressor, and the fine-tuned regressor for RNAPII. The regressor for Aldolase used an initial learning rate of 0.0005 and a batch size of 128, with the training data size of 1197 and the validation data size of 2341. The general regressor used an initial learning rate of 0.001 and a batch size of 256, with the training data size of 80443 and the validation data size of 20135. The data used to train the general regressor consisted of hole images from different grid types and hole sizes. These grids included Quantifoil 1.2/1.3 and 2/2 as well as UltrAuFoil 1.2/1.3 from samples that ranged in molecular weight from ∼50 kDa to ∼1 MDa from images in the Cianfrocco laboratory from 2017-2021. The fine-tuned regressor for RNAPII used the general regressor as the pre-trained model and only the final fully-connected layer was trainable parameters. It was trained for 20 epochs with an initial learning rate of 0.0005 and a batch size of 64. The training data size was 4544 and the validation data size was 4109.

### RL Network

We applied the deep Q-learning approach(Mnih et al., 2013) to learn a planning policy for cryo-EM data collection. The DQN used in our work is a 3-layer MLP with ReLU activations. The size of each layer is 128, 256, and 128, respectively. Since moving the microscope to any target hole is considered a unique action from the agent in our RL system, the action space of the system has the same size as the training samples, which can be potentially large. In addition, during tests, the number of holes (i.e., actions) may vary case by case. Because of this, the general practice as many approaches do, which sets the number of output nodes in the network to the number of actions, is not practical for our case. While more sophisticated methods(Lazaric et al., 2007; Le et al., 2022) exist for dealing with varying space sizes, we applied a single output node to estimate the Q-value of a hole once each time and batch process all the actions efficiently. A detailed description of the components in our RL setup and training can be found in(Fan et al., 2022). In this paper, we set the duration in our system to 480 minutes for training.

### Features to DQN

A good RL policy, if possible, should always reward small microscope movements. In other words, the hole path planning strategy should prioritize areas with rich low CTFMaxRes holes first. The number of low CTFMaxRes holes on a patch (square or grid) image is closely related to the quality (or value) of the image in cryo-EM data collection. Based on this, we design hierarchical features for the DQN that represent the quality of images at different levels, including histograms of CTFMaxRes on unvisited holes and the number of visited and unvisited low CTFMaxRes holes. The histograms are smoothed by a Gaussian kernel to improve robustness against the imperfect regression results. We listed the details of these features, which are computed separately for each hole and linearly normalized between 0 and 1 in **Extended Data Table 2**. We also considered the history of the microscope movement by concatenating the features for the last *k* − 1 visited holes and the current one to be visited to form the final input to the DQN. In this study, *k* was empirically set to 4.

### Cryo-EM simulator and human study

We conducted a human study to directly compare the performance of cryoRL with humans. The dataset used was Aldolase^Au^. We developed a simulator that mimics real data collection (**Supplemental Movie 1**) without the waiting time to acquire each image. The hierarchical layers in the dataset were preserved and well reflected in the simulator. The tested users were asked to collect 450-500 micrographs, and the resulting trajectories were recorded. We asked the users to collect as many as good micrographs (with low CTFMaxRes), while trying to minimize the time cost of moving the stage to distance positions, as in a real data collection. The users were given the freedom to look through the low magnification images at different magnification hierarchies without penalty to provide a fair competition with cryoRL. We note that this type of analysis is not possible with the traditional “sequential” data collection strategy. The tested users were given the same time cost table (**Extended Data Table 1**). We then translated the trajectories with the time costs used in this manuscript and truncated the trajectories at 480 min to compare them with the performance by cryoRL on the same dataset. The simulator has two modes, “inspection mode” and “queueing mode”. In the inspection mode, the ground truth CTFMaxRes will be given once the hole is “collected”. In the queueing mode, the user may queue up a number of hole targets without knowing their ground truth CTFMaxRes values until switching to the inspection mode. The tested users include experts who are scientists with over 8 years of cryo-EM data collection experience and are still active in collecting data, and novices who have less than 1 year of experience in cryo-EM data collection but have prior knowledge of cryo-EM data collection. No significant difference was found between the average performances of the experts and the novices.

### Cryo-EM data processing

CTF estimation was performed in Warp(Tegunov and Cramer, 2019) for the complete Aldolase^Au^ dataset (5822 total micrographs) and all trajectories, filtering from 30-3 Å and 0.2-10.0 µm defocus. Particles were picked with an expected diameter of 60 Å using WARP’s BoxNet2 neural network model (0.98 Å/pixel, 216 px box size). The remainder of the processing was conducted in CryoSPARC(Punjani et al., 2017), beginning with 2D classification (150 classes, no specified mask diameter). Particles corresponding to 2D class averages with a resolution below 10 Å were excluded from downstream processing. An initial 3D homogeneous refinement was performed on the remaining particles with C1 symmetry and EMD-21023(Wu et al., 2020) as the initial model. Particles and volumes from this initial refinement were then subjected to a 3D classification (5 classes, D2 symmetry) using CryoSPARC’s heterogeneous refinement tool. For human and cryoRL-selected trajectories, particles corresponding to the most populated 3D class were auto-refined with D2 symmetry. All resolution estimates are defined using a Fourier shell correlation (FSC) cutoff of 0.143. Tight masking yielded a final resolution of ∼4.0 Å averaged across 10 randomly sampled cryoRL trajectories and an average of ∼4.7 Å from the 10 trajectories by human test subjects, according to gold-standard FSC. Particles from the complete Aldolase^Au^ dataset were subjected to a second round of 3D classification (5 classes, D2 symmetry) before a final 3D auto-refinement with D2 symmetry. The final resolution of the complete Aldolase^Au^ dataset was ∼3.8 Å with tight masking applied. Images of all maps were captured using UCSF Chimera(Pettersen et al., 2004) (**Extended Data Fig. 5**).

### Jaccard similarity coefficient

The Jaccard similarity coefficient was calculated with the standard formula:

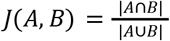

where A and B are the two sets of micrographs that two different users or cryoRL collected.

### Cryo-EM sample preparation and data collection

Aldolase was prepared using rabbit muscle aldolase (Sigma Aldrich) at 1.6 mg/ml in a buffer of 20 mM HEPES (pH 7.4) and 50 mM NaCl onto either a freshly glow discharged Quantifoil 1.2/1.3 or UltrAuFoil grid plunge frozen using a Vitrobot Mark IV (Thermo Fisher Scientific). Recombinant mouse Apoferritin was purified as described in a protocol by Dr. Christos Savva. The sample was diluted to 4.1 mg/mL in buffer 50 mM Tris (pH 7.5), 100 mM NaCl, and 500 μM TCEP prior to application on a freshly glow discharged UltrAuFoil 1.2/1.3 grid using a Vitrobot Mark VI (Thermo Fisher Scientific). RNAPII was purified from *S. cerevisiae* as described(Farnung et al., 2021, 2018). An RNAPII elongation complex was formed with factors Spt6, Spt4/5, and the Polymerase-associated complex 1 complex on a linear DNA construct as described(Vos et al., 2020). The complex was purified by size exclusion chromatography on an Äkta Micro (Cytvia) equipped with a Superose 6 3.2-300 column in a buffer containing 20 mM Na-HEPES pH 7.4, 75 mM NaCl, 3 mM MgCl2, 4% (v/v) glycerol. Fractions containing complex were assessed by SDS-PAGE, crosslinked with glutaraldehyde (PMID: 32541898), quenched, and dialyzed against a buffer containing 20 mM Na-HEPES pH 7.4, 75 mM NaCl, and 3 mM MgCl2 for 4 hrs at 4°C. The sample (final concentration 150-200 nM) was applied to freshly glow discharged Au-Flat 1.2/1.3 grids using a Vitrobot Mark VI (Thermo Fisher Scientific).

We imaged Aldolase and Apoferritin using a Talos Arctica (Thermo Fisher Scientific) equipped with a K2 Summit detector (Gatan Inc.) operating at 200 keV. Micrographs were collected in counted mode using a pixel size of 0.91Å/pixel. Aldolase^Au^ was imaged using a Glacios (Thermo Fisher Scientific) equipped with a K2 Summit detector (Gatan Inc.) operating at 200 keV with a pixel size of 0.98Å/pixel in counted mode. RNAPII was imaged using a Titan Krios G2 equipped with a K3 detector (Gatan Inc.) behind a Gatan Imaging Filter (Gatan Inc.) operating at 300 keV with a slit width of 20 eV.

All data were collected using Leginon(Suloway et al., 2005). Movies were motion-corrected using MotionCor2(Zheng et al., 2017) and CTF-estimated with CTFFIND4(Rohou and Grigorieff, 2015) run from within the Appion environment(Lander et al., 2009).

### CryoRL specifics per grid

To train and test cryoRL on Aldolase, we collected a dataset of Aldolase using a Talos Arctica microscope equipped with a K2 detector. The dataset consisted of 3538 holes from 25 squares (**Fig. 2A**) and was split into training and test sets (1197 holes originating from 9 squares belong to the training set, and 2341 holes from the remaining 16 squares belong to the test set). Note that the training set represents only a small subset of the total data (33.8%). We trained both the deep regressor and RL network on the training set and applied the models on the test set to quantify the performance.

To test the transferability of RL models, we collected datasets of Apoferritin and RNA Polymerase II (RNAPII). The specifics of the regression models training and testing can be found in the “Deep regressors” section above. Specifically, the grid type used in RNAPII was not in the training set of the general regressor, therefore the prediction of the general regressor on RNAPII was poor. After fine-tuning, the performance of the regressor was still not optimal, but a general trend could be observed (**Extended Data Fig. 2C**). For RL networks, we simply used the RL model trained on the Aldolase dataset as a transferability test. We then applied the regressors and RL network to the test set to quantify the performance.

For Aldolase^Au^, we collected the systematic dataset consisting of 5822 holes from 30 squares. The general regressor and RL network trained on Aldolase were applied on this dataset to quantify the performance and compare it with the human study.

## Extended Figures

**Extended Data Figure 1.**
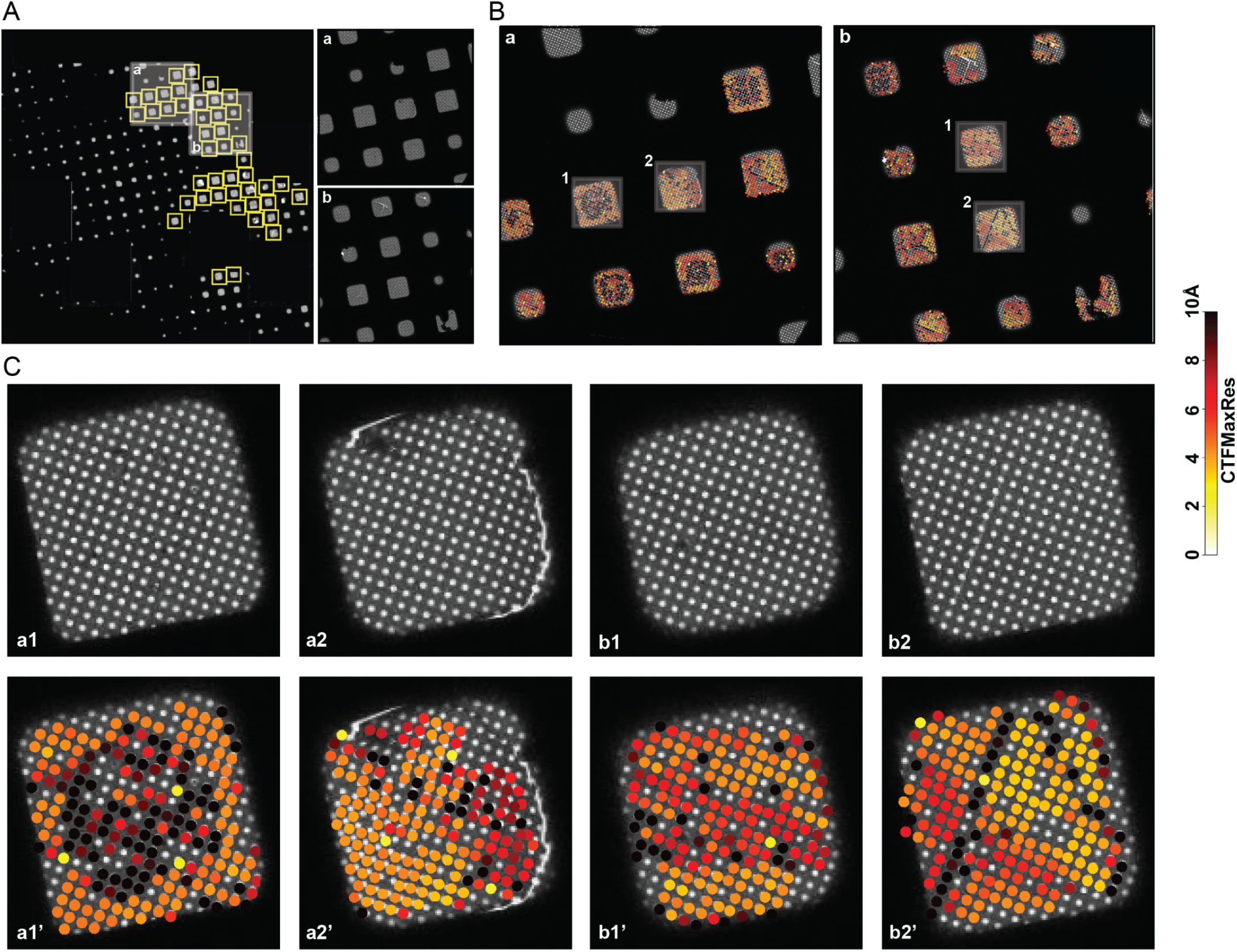
Cryo-EM grids are complicated data landscapes. (A) Overview of cryo-EM grid used for systematic data collection. Yellow squares indicate regions where almost every hole was collected. Regions a and b are shaded and zoomed in. (B) Zoom-in on the shaded regions a and b in (A), with ground truth CTFMaxRes values overlayed. (C) Zoom-in on the selected squares in (B) (top), with ground truth CTFMaxRes values overlaid (bottom). (B) and (C) share the same color bar. Note that in real data collection, ground truth CTFMaxRes values are unknown before the holes are visited.

**Extended Data Figure 2.**
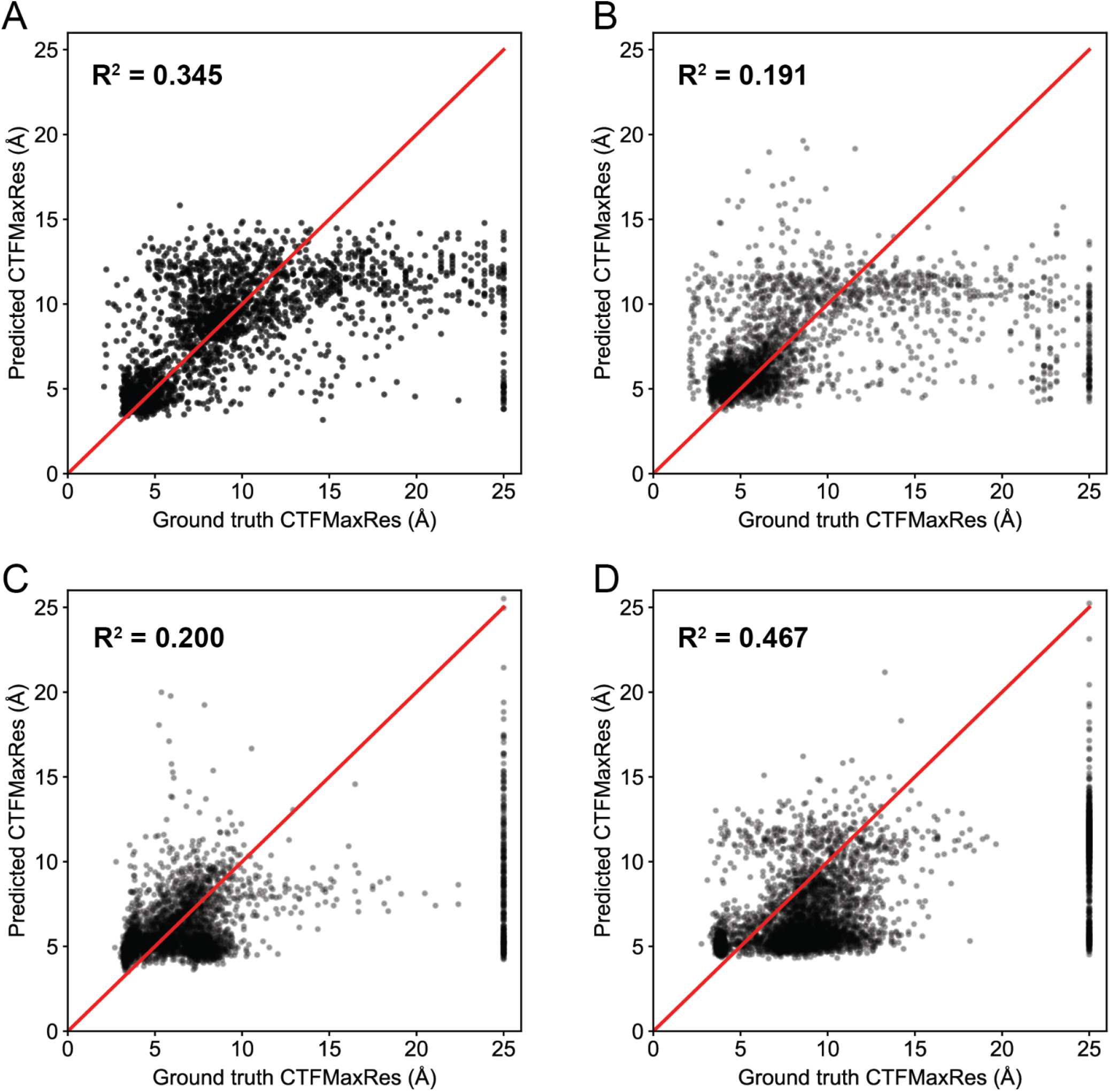
Performance of the regressors on the test datasets. (A) Performance of the regressor on the Aldolase dataset. The regressor was trained on a subset of the Aldolase dataset and tested on the rest of the dataset. (B) Performance of the general regressor on the Apoferritin dataset. (C) Performance of the regressor (fine-tuned from the general regressor) on the RNAPII dataset. (D) Performance of the general regressor on the Aldolase^Au^ dataset.

**Extended Data Figure 3.**
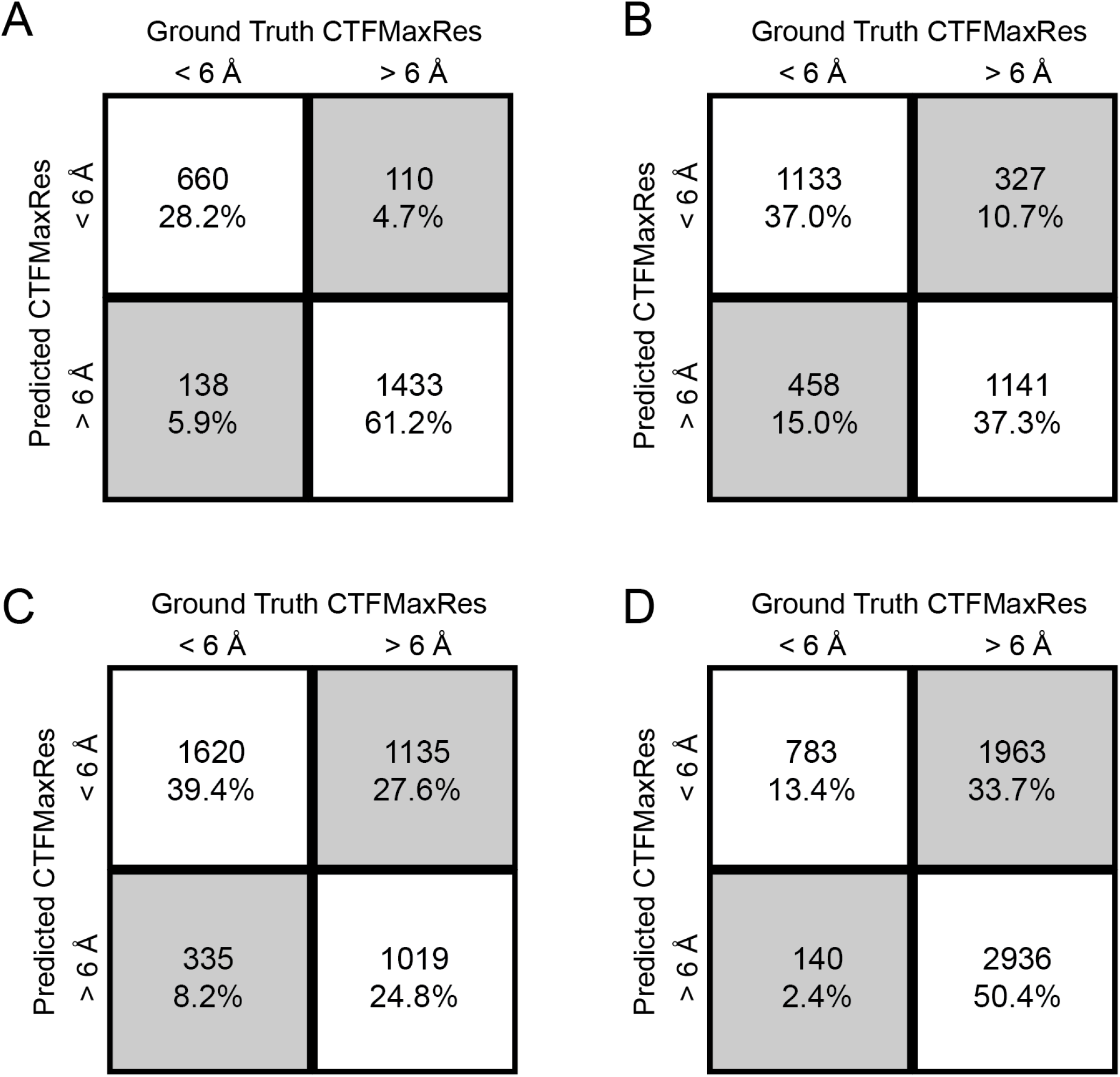
Confusion matrices of the regressors on the test datasets. (A) Confusion matrix of the regressor result on the Aldolase dataset at a cutoff of 6 Å. The regressor was trained on a subset of the Aldolase dataset and tested on the rest of the dataset. (B) Confusion matrix of the general regressor result on the Apoferritin dataset at a cutoff of 6 Å. (C) Confusion matrix of the regressor (fine-tuned from the general regressor) result on the RNAPII dataset at a cutoff of 6 Å. (D) Confusion matrix of the general regressor

**Extended Data Figure 4.**
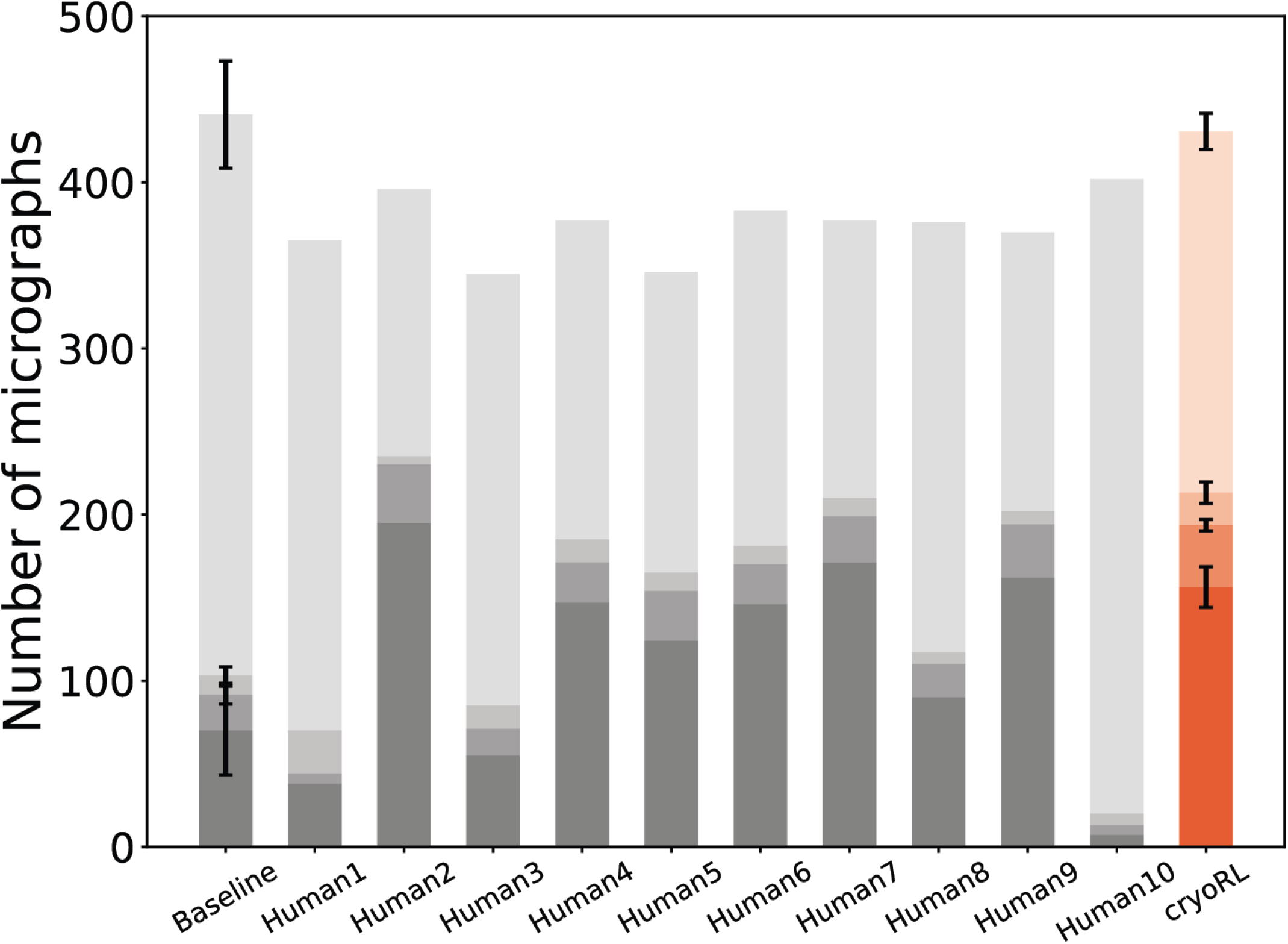
Performance of each human test subject on the Aldolase^Au^ dataset. Detailed performance of each human test subject in the human study, compared with the performance of the baseline and cryoRL. Different grayscale colors indicate CTFMaxRes below 4 Å, between 4 Å and 5 Å, between 5 Å and 6 Å, and above 6 Å, as in the main text figures.

**Extended Data Figure 5.**
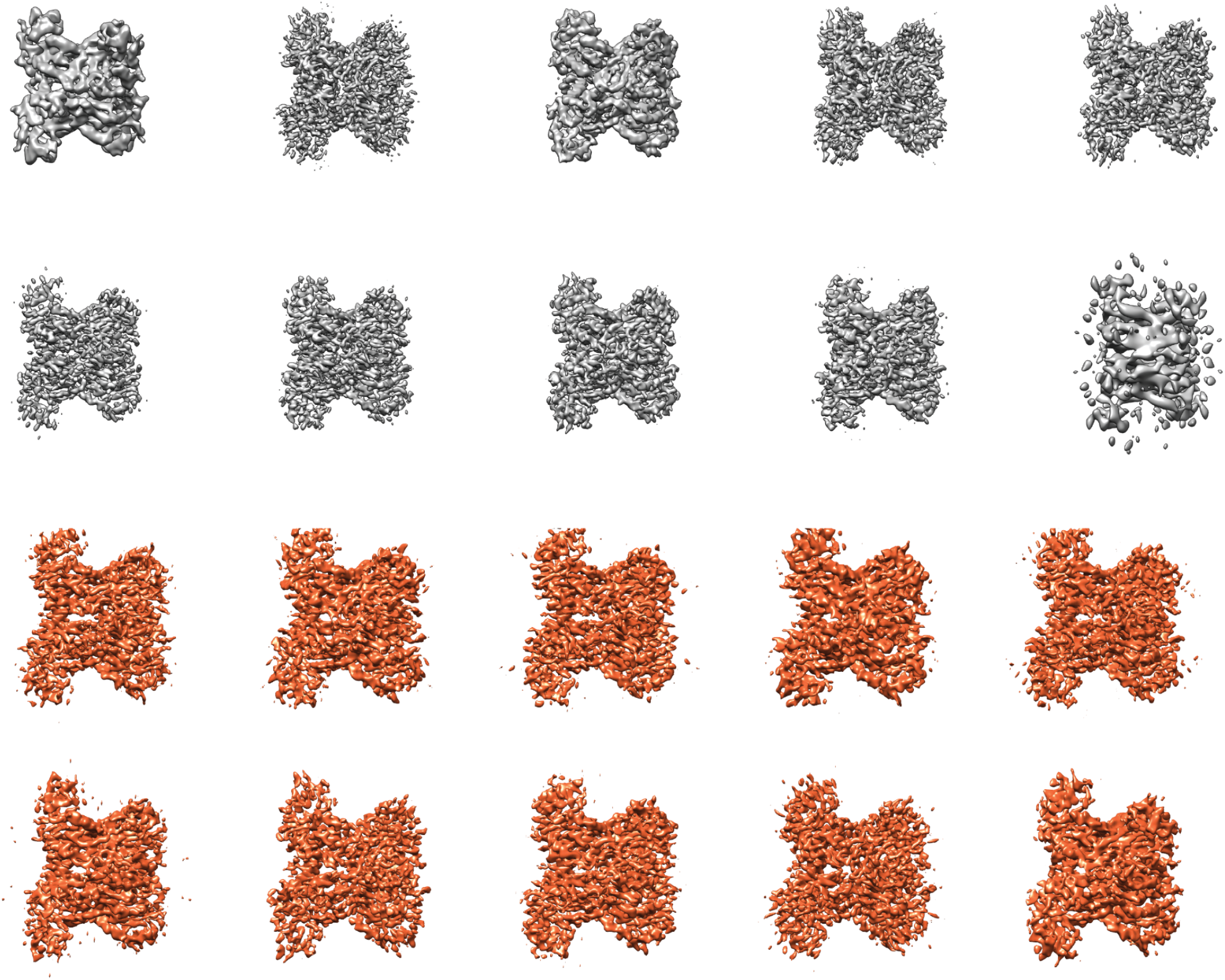
Sharpened 3D reconstructions of Aldolase^Au^ from human and cryoRL trajectories. Reconstructions from human subjects (gray) and cryoRL (orange).

**Extended Data Table 1.**
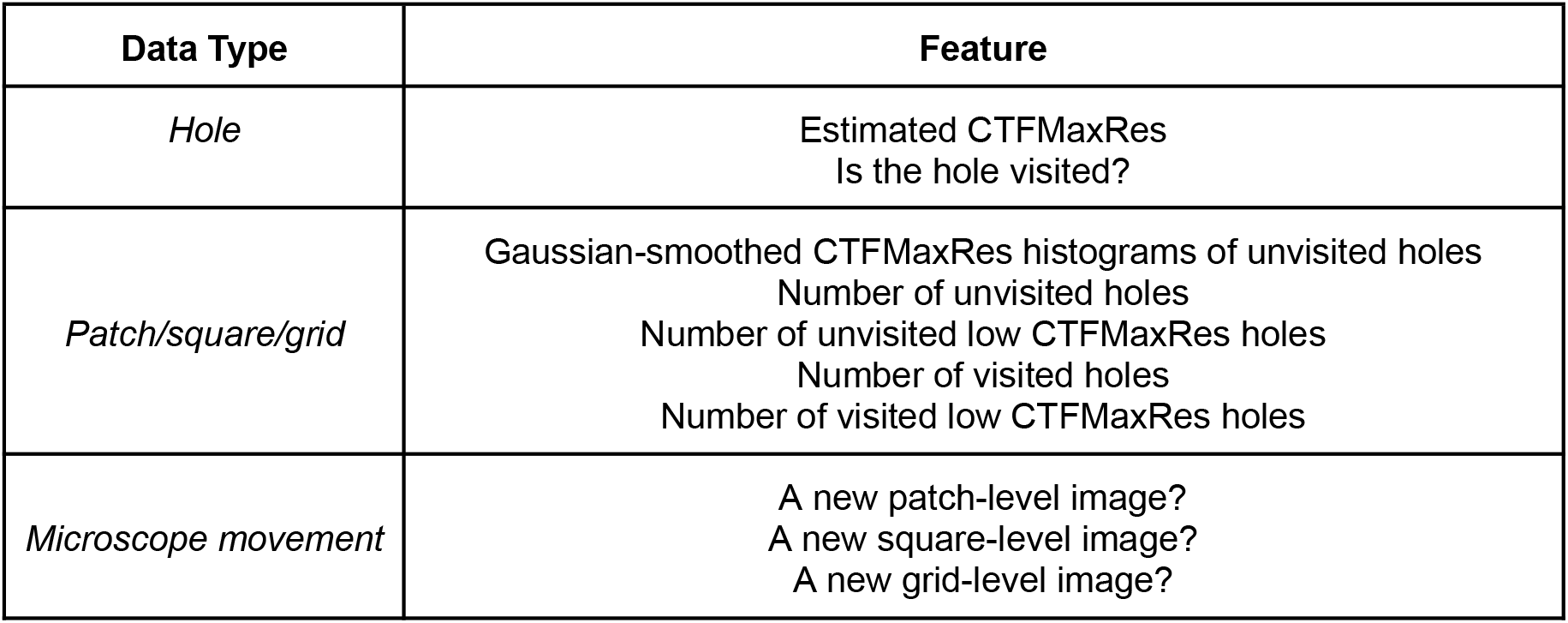
Input features to the deep-Q network

| **Data Type** | **Feature** |
| --- | --- |
| *Hole* | Estimated CTFMaxRes<br>Is the hole visited? |
| *Patch/square/grid* | Gaussian-smoothed CTFMaxRes histograms of unvisited holes<br>Number of unvisited holes<br>Number of unvisited low CTFMaxRes holes<br>Number of visited holes<br>Number of visited low CTFMaxRes holes |
| *Microscope movement* | A new patch-level image?<br>A new square-level image?<br>A new grid-level image? |

**Extended Data Table 2.**
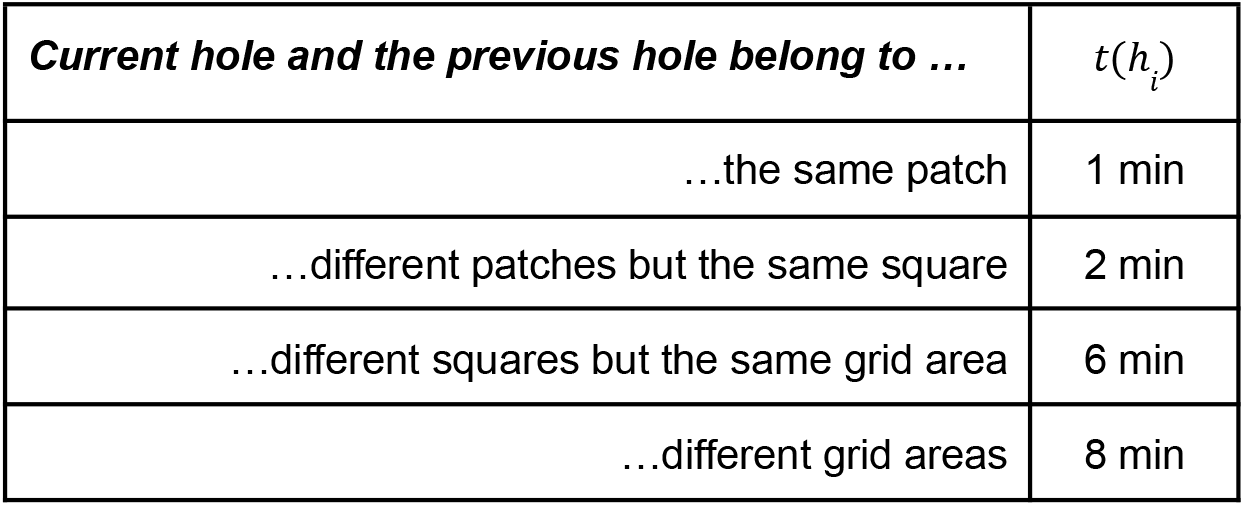
Time cost table used in this study.

**Extended Data Table 3.**
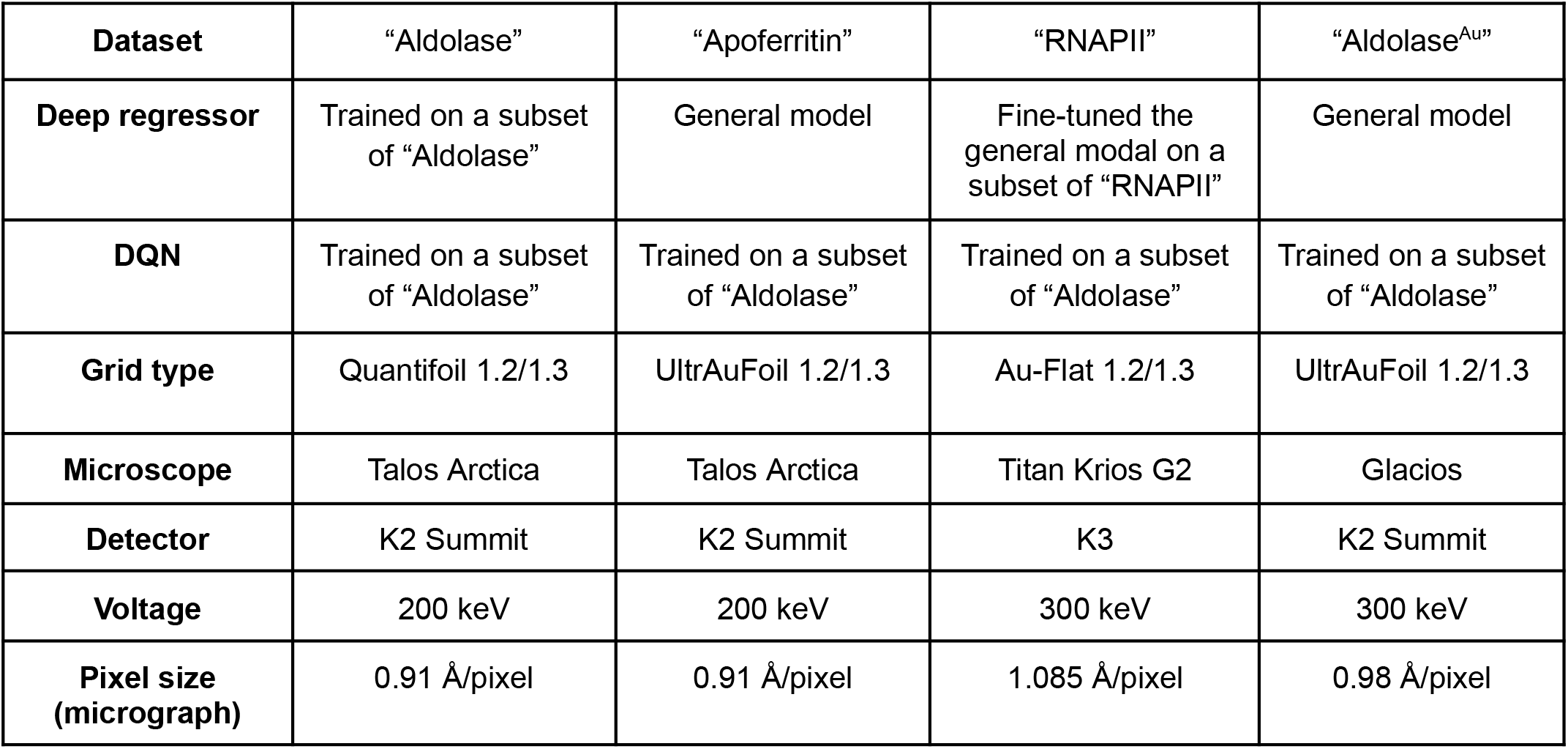
Detailed information of the four experiments presented in this paper.

## Supplemental Data

Supplemental Movie 1 - Demonstration of the cryo-EM data collection simulator used in the human study.

